# Probiotics for the prevention of antibiotic-associated adverse events in children – a systematic review to inform development of a core outcome set

**DOI:** 10.1101/2020.01.27.920892

**Authors:** Jan Łukasik, Qin Guo, Leah Boulos, Hania Szajewska, Bradley C. Johnston

**Affiliations:** Medical University of Warsaw, Department of Pediatrics, Warsaw, Poland; West China Second University Hospital, Department of Pediatrics, Chengdu, China; Maritime SPOR SUPPORT Unit, Halifax, Canada; Dalhousie University, Department of Community Health and Epidemiology, Halifax, Canada

## Abstract

**Introduction:** Routine use of probiotics during antibiotic therapy in children remains a subject of discussion. To facilitate synthesis of individual study results and guideline formulation, it is important to assess predefined, similar, and clinically important outcomes. Core outcome sets are a proposed solution for this issue. Aim of this review was to document choice, design, and heterogeneity of outcomes in studies that assessed the effects of probiotics used for the prevention of antibiotic-associated adverse events in children.

**Methods:** A systematic literature search covering three major databases was performed. Trials that evaluated oral probiotics’ use concomitant with antibiotic therapy in children were included. Data on outcome definitions, measurement instruments, and follow-up were extracted. The outcomes were assigned to predefined core areas and domains. Data were analyzed descriptively.

**Results:** Thirty-six trials were included in this review. Diarrhea, the most commonly reported outcome, had diagnostic criteria clearly defined only in 20 trials. In total, sixteen different definitions of diarrhea were identified. Diarrhea duration, severity and etiology were reported in 8, 4 and 6 studies, respectively. Nineteen studies assessed gastrointestinal symptoms other than diarrhea. Seven studies reported outcomes related to resource use or the economic impact of the intervention. Only 2 studies assessed outcomes related to life impact. None of the studies predefined adverse events of probiotic use.

**Conclusions:** Identified outcomes were characterized by substantial heterogeneity. Majority of outcomes were not designed to evaluate endpoints of real-life relevance. Results from this review suggest the need for a new core outcome set consisting of outcomes important for decision-making.

## Introduction

The human gastrointestinal tract is colonized by hundreds of different microorganisms, which together form the gut microbiota (1, 2). Use of antibiotics is one of the factors known to alter the microbiota composition, which in turn may have an effect on an individual’s health. Typical adverse events associated with antibiotic use include various gastrointestinal symptoms such as diarrhea, nausea, vomiting, and abdominal pain (3). Among them, antibiotic-associated diarrhea (AAD), often defined as ‘diarrhea that occurs in relation to antibiotic treatment with the exclusion of other etiologies’ (4), is the best documented.

Over 30 randomized controlled trials (RCTs), mostly with probiotics as an intervention, have been performed to assess the prophylactic strategies for AAD in children (5). In the largest observational study of 650 children published in 2003, the estimated AAD incidence in the pediatric outpatient population was 11% (6). On the other hand, in a recent (2019) Cochrane review (5), the incidence of AAD varied greatly from study to study, ranging from 2% (7) to 80% (8). In addition to estimates sometimes being derived from very small underpowered studies (8–11), one of the factors responsible for this heterogeneity in reported incidences could be the definition of AAD adopted by authors of different RCTs and the methods used for measurement of this outcome. Among others, AAD diagnostic criteria vary between the studies in the terms of stool frequency, time from the start of antibiotic therapy, and microbiological methods, if any, used to exclude other etiologies of diarrhea.

Other potential effects of early-life microbiota alterations include later-life consequences such as obesity (12), allergies (13), autoimmune disorders (14), and neurodevelopmental abnormalities (15). The long-term health impact of probiotics and antibiotics administered during infancy has been evaluated in some RCTs (16, 17), but this outcome is not a part of a routine trial design.

According to the 2016 European Society for Pediatric Gastroenterology, Hepatology, and Nutrition (ESPGHAN) guidelines, some probiotic strains may be effective in AAD prevention (4). Consistent with this, a 2019 Cochrane systematic review of 33 studies concluded that there is a moderate protective effect of probiotics for preventing AAD (5). Still, this use of probiotics is the subject of a lasting discussion due to their cost, and the fact that AAD is usually a mild and self-limiting disease (18). To draw practical conclusions from RCTs, it is important to assess AAD severity and its impact on the patient’s everyday life, including global assessment and health-related quality of life, with agreed-upon definitions and outcomes. However, a 2010 systematic review of outcomes used in trials of pediatric acute diarrhea revealed substantial heterogeneity in both the definitions of and the measurement methods for diarrhea (19). Similarly, in the 2019 Cochrane systematic review, the criteria for defining the incidence of diarrhea according to each primary investigator’s definition varied widely among the studies (5). Differences in reported definitions, outcomes, and their measurement methods between studies may lead to difficulties in synthesizing results and hinder the process of guideline formulation. Standard definitions for main outcomes are a possible solution to these issues, and systematic reviews addressing the choice of outcomes in already performed studies are one of the first steps in the process of designing a core outcome set (COS) (20). In 2016, a document by the Consensus Group on Outcome Measures Made in Pediatric Enteral Nutrition Clinical Trials (COMMENT) was published, proposing core outcomes for future use in RCTs evaluating therapeutic and preventive strategies for acute gastroenteritis (21). However, authors of this document did not include any statements regarding outcomes specific for AAD. Also, no core outcome set to date has been proposed for use in trials in which probiotics are administered concurrently with antibiotic therapy.

Our primary aim was to systematically document the definitions of AAD, as well as all of the methods used to measure and describe this outcome, in studies that assessed the effect(s) of probiotics used for AAD prevention. Additionally, we aimed to document any other outcomes reported in studies on probiotic use during antibiotic therapy, provided that they were used to examine probiotics’ effect(s) in the prevention of antibiotic-associated adverse events.

## Methods

### Inclusion/Exclusion Criteria for the Review

Studies that evaluated oral probiotics’ potential to prevent adverse events associated with antibiotic therapy were eligible for inclusion in this review. Eligible studies could be RCTs, non-randomized trials (NRTs), or observational studies (e.g., cohort studies, case-control studies) and had to be conducted in a population of children up to 18 years of age. Among the studies conducted in mixed populations of children and adults, only those that reported separate data for a subgroup of children were included. Furthermore, only studies published in English were included.

Studies that reported only laboratory outcomes (e.g., only stool microbiota composition) were not included in this review. Since the main focus of this review was the prevention of AAD, studies on probiotics used concurrently with antibiotics in the treatment of *Clostridium difficile*-associated diarrhea or other types of diarrhea were excluded. Additionally, studies conducted exclusively in premature infants and in critically ill children hospitalized in intensive care units were also not included, because the characteristics of these populations and the goals of probiotic use differ greatly from those in the general population.

### Search methods

A systematic search was performed from inception to October 23, 2018 in three major databases (MEDLINE, Embase, and CENTRAL). The search strategy was developed by an information specialist and included controlled vocabulary and keywords related to ‘antibiotic’ and ‘probiotic’ terms. The full search strategy for the MEDLINE database is available in S1 Table. Additionally, references of relevant review articles were manually searched.

### Selection of studies

JŁ screened the search results and identified abstracts of potentially eligible articles. After abstract screening, full articles were acquired and independently evaluated for eligibility by JŁ and QG. Any disagreements concerning eligibility were resolved by discussion between the authors, and if needed, with a senior researcher.

### Data extraction

The data from the included studies were extracted using an abstraction form developed specifically for this review. Extracted data included standard characteristics of studies (author, publication year, country, study type and setting, age and number of participants, indication for antibiotic treatment, type of antibiotics, investigated probiotic, and type of control group) and data specific to the outcomes. Each identified outcome was assigned to one of 4 core areas: “life impact”, “resource use”, “pathophysiological manifestations” or “death”, in accordance with the OMERACT Filter 2.0 (22). Specific outcomes were also assigned to one of the predefined outcome domains included within the core areas. In case of identification of an outcome not falling into any of the predefined domains, a new domain was created. An explanation of the outcome-related taxonomy used in the article is presented in Table 1. The data extraction and assignment of the outcomes to the core areas and domains were done independently by JŁ and QG, and any differences in opinion were resolved by discussion. The data extracted for each identified outcome included: outcome name in accordance with the terminology used in the original publication, outcome characteristics (e.g., incidence, duration, severity, primary/secondary outcome), outcome definition, outcome measurement instruments, and follow-up. The outcome was considered as primary if either: 1) the authors of the original study declared it as such, or 2) a sample size calculation was performed for this specific outcome. The data for purely biochemical or microbiological outcomes (e.g., microbiota composition) were not extracted, because their documentation and evaluation would require an entirely different methodological approach.

**Table 1.** Definitions of the terminology used in the article, in accordance with OMERACT definitions^22^

| Term | Definition | Examples |
| --- | --- | --- |
| Core area | An aspect of health or a health condition that needs to be measured to appropriately assess the effects of a health intervention. Core Areas are broad concepts consisting of a number of more specific concepts called domains. | Pathophysiological manifestations, life impact, resource use/economic impact |
| Outcome domain | An aspect of the effect of illness, categorized within the core area, but still relatively broad. | Diarrhea, gastrointestinal symptoms, absenteeism, need for additional medical procedures. |
| Outcome | Any identified result in a domain arising from exposure to a causal factor or a health intervention. | Diarrhea incidence, number of school absence days, need for intravenous rehydration. |
| Outcome measurement | A tool chosen to assess the outcome. | Visual stool form scale, symptom questionnaire, |

|  |  |  |
| --- | --- | --- |
| instrument |  | immunoassay tests for rotavirus detection. |

### Assessment of risk of bias in the included studies

Risk of bias (RoB) assessment is not a mandatory part of systematic reviews of outcomes (20); however, we decided to present it for informative purposes. The Cochrane Collaboration’s Tool for Assessing Risk of Bias (23) was used for RCTs and non-randomized trials. Wherever possible, we present the RoB assessment derived from the recent Cochrane review (5). For the remaining studies, the RoB assessment was performed by JŁ.

### Data analysis

Data on the identified outcomes are presented in numbers and percentages and analysed descriptively. Since this review aims to document the methods of outcome measurement and reporting, no analysis of the treatment effects was performed.

## Results

### Search results and overall characteristics

In total, we identified 4251 records by the database search and additional 369 records from the review articles’ references. After exclusion of duplicates and title and abstract screening, full texts of 80 articles were assessed for eligibility. After full-text assessment, 36 articles ultimately met the inclusion criteria for this review(7–11, 24–54). The flow diagram of the study selection process is presented in Fig 1. Reasons for exclusion of the specific studies are presented in S2 Table.

**Fig 1.**
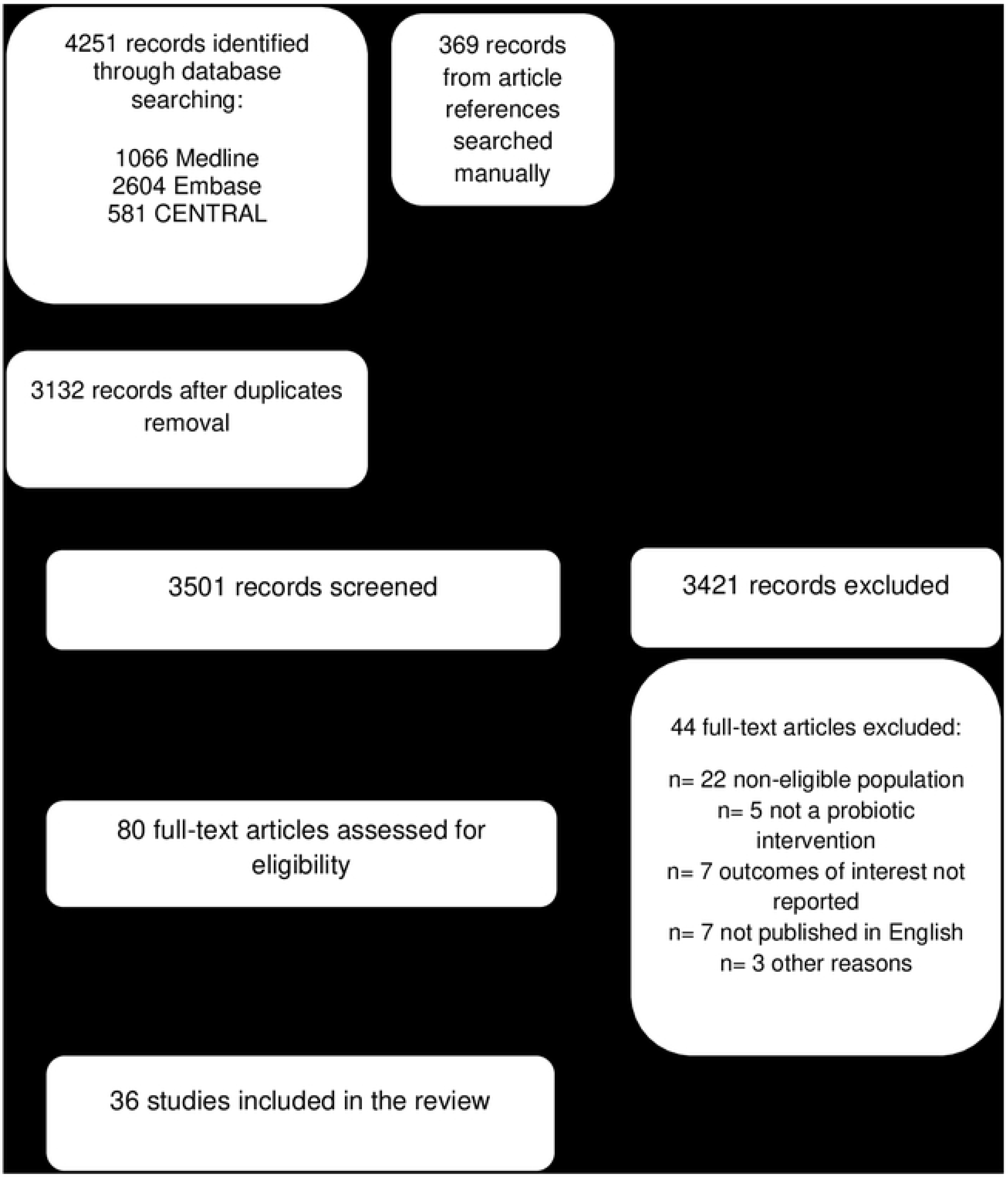
Flow chart diagram

Among the included studies, 32 (89%) were RCTs, and the remaining 4 were NRTs. The total number of participants was 5506, ranging from 18 to 653 children in individual trials. Ten trials were conducted in the inpatient setting, 14 in the outpatient setting, 5 in the mixed setting, and 1 in an unclear setting. Additionally, in 6 trials on *H. pylori* treatment, the setting was not clearly defined; however, we assumed it to be ‘probably outpatient’, as *H. pylori* eradication usually takes place at home. The most common indications for antibiotic therapy were *H. pylori* treatment (11 studies, 31%), various childhood infections (11 studies, 31%), and respiratory tract infections (7 studies, 19%). Various beta-lactams were most often used (31 studies, 86%), followed by macrolides (22 studies, 61%). The majority of the trials (19 studies, 53%) used single-strain probiotics as an intervention and were placebo-controlled (21 studies, 58%). A summary of the included studies’ characteristics is presented in S3 Table. All of the identified outcomes and their characteristics are presented in Tables S4 & S5.

The RoB in the included trials varied. Most of the studies were characterized by substantial RoB. A summary of the RoB assessment is presented in S1 Fig.

### Outcome domain: diarrhea

The occurrence/incidence of diarrhea was reported as an outcome in 32 (89%) of the included studies and 20 (63%) of these studies reported it as a primary outcome. In only 20 (63%) of these 32 studies were the criteria for diarrhea diagnosis clearly defined. In the remaining studies, the occurrence of diarrhea was reported by parents or patients during interviews or in study diaries, and diagnosed based on the participants’ or investigators’ judgment, with unclear diagnostic criteria. In 9 (28%) of the studies which assessed this outcome, various stool form scales were used, most commonly (7 studies) the Bristol Stool Form Scale (BSFS) (55).

Based on the frequency and minimal duration of loose stools occurrence, 8 different definitions of diarrhea were used by the authors of the original studies. Most commonly (11 studies, 31%), diarrhea was diagnosed when at least 3 stools of abnormally loose consistency occurred during 48 hours. However, when different definitions of “abnormal stool consistency” were taken into an account, as many as 16 different definitions of diarrhea were identified. The most commonly used definitions of diarrhea are presented in Fig 2.

**Fig 2.**
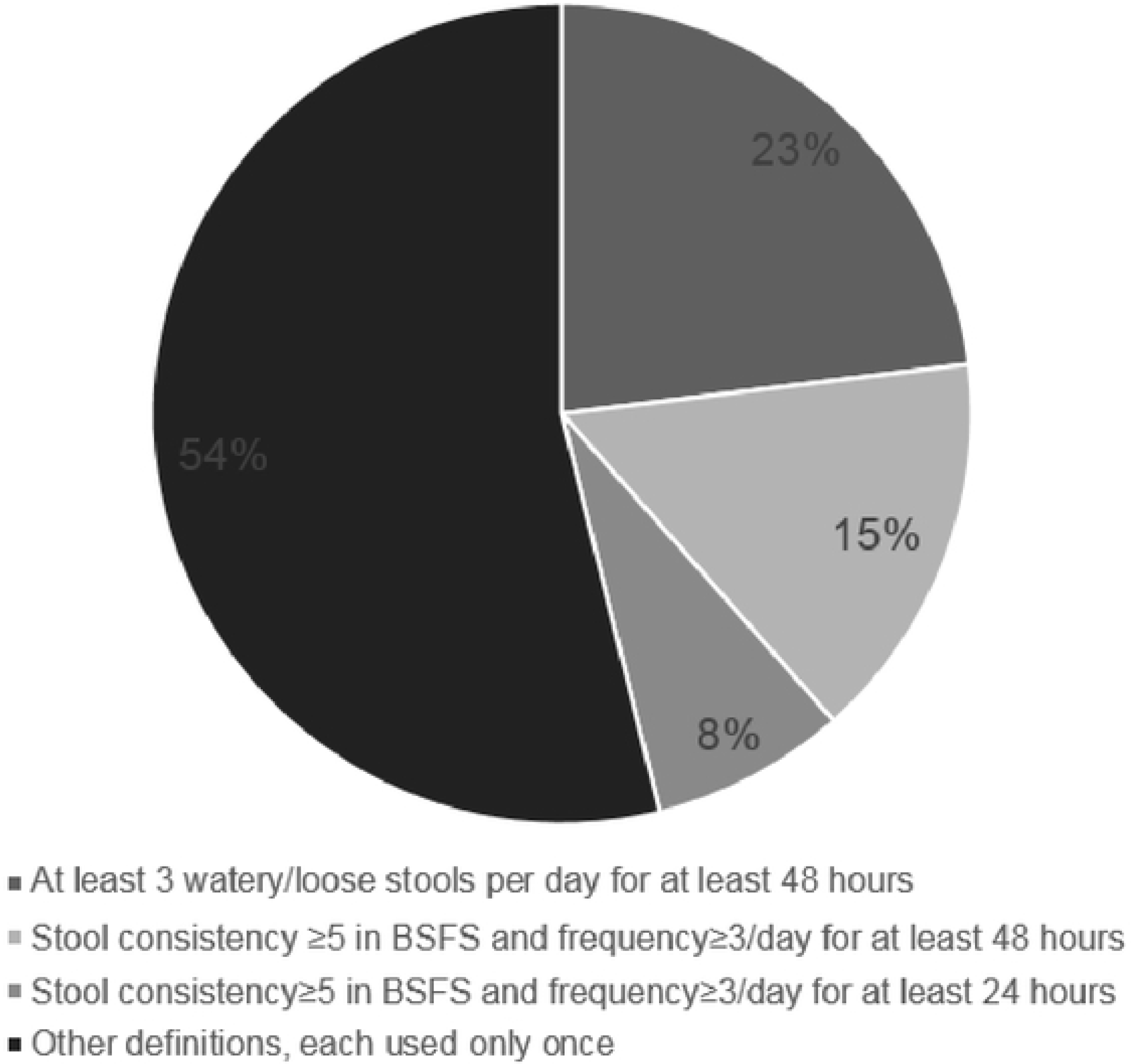
Most commonly used definitions of diarrhea.

Surprisingly, among the 32 studies that reported data on diarrhea occurrence, the authors referred to their outcome as ‘antibiotic-associated diarrhea’ or ‘treatment-associated diarrhea’ in only 13 articles (39%). Among them, only 6 studies (19%) investigated a potentially infectious origin of diarrhea. Moreover, in 2 of them, the authors did not utilize this information to support or exclude a diagnosis of AAD (9, 29). Authors of the other 4 studies diagnosed AAD as “diarrhea caused by *C. difficile* or of otherwise unknown origin” and performed enzyme immunoassay tests for rota- and adenoviruses detection and stool cultures for bacterial pathogens (35, 37, 42, 45). A single study additionally tested for norovirus infection using enzyme immunoassay (35).

Included studies varied with respect to follow-up duration. In 21 (66%) of the 32 trials that assessed diarrhea as an outcome, the incidence of diarrhea was assessed during antibiotic treatment and an additional follow-up period, which varied from 1 week after the end of antibiotic therapy (32, 39) to up to 7 months after its beginning (36). Seven studies (22%) assessed diarrhea only during antibiotic treatment (28, 30, 31, 35, 50, 51, 54), and 3 studies (9%), only during the first 3 to 6 days of antibiotic therapy (27, 41, 43).

Among other characteristics of the diarrhea, its duration was reported in only 8 out of 32 studies, which corresponds to 25% of the studies with diarrhea as an outcome. In 5 of these studies, the duration was not defined (8, 26, 27, 29, 51), whereas in each of the 3 remaining studies its definition varied (28, 31, 45). Diarrhea severity was reported as an outcome in only 4 of the studies (13%), and it was defined differently in every one of them, usually on the basis of discharge frequency and stool consistency (7, 26, 28, 32). Diarrhea duration and severity were reported as co-primary outcomes in one study each (32, 45), while in the other studies they were either secondary or unspecified outcomes. Where provided, the definitions of diarrhea duration and severity can be found in S5 Table.

Other outcomes regarding diarrhea included occurrence of infectious diarrhea - 5 studies (26, 29, 35, 37, 42), stool consistency regardless of diarrhea occurrence - 5 studies (31, 38, 39, 51, 52), bowel movement frequency - 3 studies (48, 51, 53), and time to diarrhea onset from the start of antibiotic therapy - 4 studies (8, 28, 29, 32). Additionally, the efficacy of diarrhea treatment, diarrhea-associated dehydration and time to first occurrence of loose stool were reported in one study each (28, 29, 32).

### Outcome domain: *Clostridium difficile* infection

In 6 studies, patients were investigated for the *Clostridium difficile* infection. In 1 study, the tests for toxin A and B were performed regardless of whether or not diarrhea occurred (i.e., asymptomatic carrier)(7), while in the other 5 they were performed only in case of diarrhea (26, 35, 37, 42, 45). One study used both the immunoassay for *C. difficile* toxin A detection and stool culture (26), whereas the others utilized only the toxin A and B immunoassays.

### Outcome domain: other gastrointestinal symptoms

The most commonly reported gastrointestinal outcomes other than diarrhea included the following: abdominal pain (15 studies, 42%), vomiting (16 studies, 44%), nausea (11 studies, 31%), lack of appetite (7 studies, 20%), constipation (9 studies, 26%), bloating (7 studies, 19%), taste problems (5 studies, 14%), and flatulence (7 studies, 19%). Other less commonly assessed outcomes included belching, abdominal discomfort, symptoms included in the Gastrointestinal Symptom Rating Score (GSRS)(56) (heartburn, acid regurgitation, sucking sensations in the stomach, borborygmus, abdominal distension, eructation, passage of stools, loose stools, hard stool, urgent need for defecation and feeling of incomplete defecation), and undefined ‘gastrointestinal complications’.

In 2 studies, (7, 10), the GSRS was used to assess the gastrointestinal symptoms (56). Additionally, a visual analog scale for abdominal pain intensity was used in one study (51), and a 3-point GI symptom rating scale was used in another (44). In the remaining studies, the gastrointestinal symptoms were reported by parents and/or children during interviews or in study diaries.

### Other outcomes from “pathophysiological manifestations” core area

None of the included studies assessed long-term adverse events associated with antibiotic use. Among the included studies, 18 (50%) reported data on adverse events potentially associated with probiotic use. In none of those studies were the adverse events predefined by the authors.

### Outcomes from other core areas

Seven studies (19%) reported outcomes from the “resource use/economical impact” core area (27, 31, 35, 37, 42, 47, 48). The most common outcomes from this area were need for antibiotic discontinuation due to diarrhea (6 studies), need for intravenous rehydration (5 studies), and need for hospitalization due to diarrhea (5 studies).

Only 2 studies assessed outcomes from “life impact” core area. A single study reported data on absence from school/day care, missed parental days at work, and overall health (38), and another study reported the data on duration of hospital stay (31).

## Discussion

In this review of outcomes used in studies assessing probiotic prophylactic interventions during antibiotic therapy in children, 32 RCTs and 4 NRTs were included. The incidence (occurrence) of diarrhea was the most commonly reported outcome. However, diagnostic criteria for diarrhea were clearly defined in only 63% of the 32 studies reporting this outcome. The majority of those studies did not utilize a validated instrument to assess the construct of diarrhea, the combination of stool frequency and consistency, did not report data on diarrhea duration and/or severity, and did not perform any microbiological tests to rule out its infectious origin. Sixteen different definitions of diarrhea were identified ranging from 1 or more abnormally loose stools per day (49) to 3 abnormally loose or liquid stools per 48 hours (9, 26, 29, 37, 42, 47, 48). The follow-up duration in the included studies also varied. Diarrhea duration and severity were often not reported, and their definitions, if provided, were different in each study. Less than half of the included studies reported data on other GI symptoms, such as abdominal pain or vomiting, and in most of them authors did not report use of any assessment instruments besides study diaries. Finally, studies rarely included outcomes from ‘pragmatic’ core areas, i.e., ‘life impact’ and ‘resource use and economical impact’.

To our knowledge, this is the first systematic review documenting the outcome measurement and reporting methods used in studies on this particular subject. Its methodology adhered both to the Cochrane Collaboration’s guidelines for systematic reviews(23) and to the recommendations of COMET (Core Outcome Measures in Effectiveness Trials) Initiative(20). Authors of this review have previous experience in probiotic and AAD research as well as in the field of systematic reviews. The potential limitations of this review result from the possibility of not including all relevant studies, since the search was limited to the articles published in English and only a basic search of the grey literature was performed (i.e., manual search within the article references). However, this review aims to document the outcomes and their definitions rather than the effectiveness of interventions. Not including all of the available studies is unlikely to influence the overall conclusions, particularly given our study team also has expertise in general pediatrics, including ongoing commitments to patient care. The other limitation of this review is lack of microbiota composition-related outcomes. The authors recognize microbiome analysis as an important element of studies on probiotics and antibiotics alike, however documentation and comparative assessment of the analysis methods requires a wholly different approach compared to clinical outcomes(57). Another important group of microbiological outcomes which is absent in this review is the antibiotic resistance(58), as none of the otherwise eligible studies reported this outcome.

Results of this review reveal substantial heterogeneity in the definitions of reported diarrhea-related outcomes. In 37% of the 32 included studies that reported the incidence of diarrhea as an outcome, the authors did not define criteria for diarrhea diagnosis, which increases the risk of reporting bias(59, 60). In the remaining studies, including the papers published subsequent to the core outcome set for use in clinical trials of pediatric acute diarrhea(21), multiple definitions of diarrhea were identified. The definitions of diarrhea duration and severity also varied. This heterogeneity may theoretically lead to difficulty in combining data from different studies for the purpose of meta-analysis(61). In the recent Cochrane review on pediatric AAD, substantial heterogeneity (I^2^=57%) was found in the analysis of diarrhea incidence (5). When subgroup analysis was based on only one definition of diarrhea (i.e., 3 or more loose/water/liquid stools per day for at least 2 consecutive days), the heterogeneity was significantly reduced (I^2^ = 15%). On the other hand, a test for interaction by diarrhea definition was not statistically significant, which suggests that different definitions of diarrhea were not the main reason for the overall heterogeneity of the result in the aforementioned review(5).

The other finding of our review concerns the criteria for AAD diagnosis. Even though the included studies investigated symptoms related to antibiotic use, authors referred to their outcome as ‘antibiotic-associated diarrhea’ in only 39% of the articles that reported the incidence of diarrhea. Moreover, infectious origin of diarrhea was investigated by microbiological methods in only 6 (19%) of studies. Considering the fact that most of the studies’ participants were either inpatients or visited healthcare facilities at the beginning of trial, they were at risk of nosocomial diarrhea(62). Not ruling out the possibility of infectious gastroenteritis in this group of patients introduces a risk of outcome misclassification. Even in studies that utilized microbiological methods to identify diarrhea etiology, it is impossible to completely rule out its infectious origin, due to the limited diagnostic accuracy of enzyme immunoassay methods(63, 64). Diarrhea reported as an outcome in the few studies which performed the microbiological testing is much more likely to be an actual AAD.

The most commonly assessed outcome from the ‘diarrhea’ domain was incidence data. Surprisingly, other outcomes that are arguably more patient important, such as diarrhea duration or severity, were rarely reported. Furthermore, even the most anticipatory criterion for diarrhea diagnosis was ‘at least 3 loose or watery stools per day for at least 48 hours’. This constitutes a relatively mild course of illness, especially assuming that the symptoms are likely to resolve on the third day after occurrence(65). Based only on the data for diarrhea incidence, it is difficult to assess whether the reported effect of any intervention was of actual importance to the patients. Other GI outcomes that could contribute to drawing clinically significant conclusions such as abdominal pain or vomiting, were only assessed in a small portion of the studies, even though they are likely to occur during antibiotic treatment(3). When they were reported, authors typically assessed incidence rather than duration or severity, again focusing on outcomes they may be less patient-important. Outcomes from ‘resource use’ and ‘life impact’ core areas, which reflect the pragmatic approach to clinical trial design, were rarely reported. The lack of available outcomes on life impact, particularly quality of life, is concerning. Although quality of life measures are not often an outcome employed in clinical trials assessing acute outcomes, there are examples in acute gastroenteritis(66). Although we did not find validated disease specific quality of life outcomes used in our target population, individualized quality of life instruments such as Measure Yourself Medical Outcome Profile (MYMOP) should be considered as a part of core outcomes(67).

The included studies also varied in the terms of follow-up duration with the majority of the studies following patients during the entire duration of antibiotic therapy and for at least one week after antibiotic cessation. Considering the usually short incubation time of AAD(68), these lengths of follow-up should be sufficient to identify most of the cases.

None of the included studies predefined outcomes from the domain ‘adverse events of the probiotic use’. This may result from the fact that the probiotics are unlikely to cause adverse events in immunocompetent children(69). Nevertheless, a clear and carefully planned documentation of adverse events is still important(70), as claims of harmful effects of probiotic use, particularly in immunocompromised patients, are being occasionally published (71).

## Conclusions

Outcomes reported in studies on probiotic use in children receiving antibiotic therapy are characterized by substantial heterogeneity. In the majority of trials, the outcomes and outcome measures are not designed to evaluate outcomes of real-life relevance such as patient and parent reported quality of life. Results from this review suggest the need for a new core outcome set with endpoints that cover the span of domains and outcomes important to patients, families and clinicians for decision-making.

## Supporting information

**S1 Fig. Risk of bias summary for the included studies**

**S1 Table. MEDLINE Search Strategy (Ovid MEDLINE(R) and Epub Ahead of Print, In-Process & Other Non-Indexed Citations, Daily and Versions(R))**

**S2 Table. Excluded studies with reasons of exclusion**

**S3 Table. Characteristics of the included studies**

**S4 Table. Outcomes identified in the included studies.**

**S5 Table. Characteristics of the identified outcomes.**

